# Forgot what you like? Evidence for hippocampal dependence of value-based decisions

**DOI:** 10.1101/170969

**Authors:** A. Z. Enkavi, B. Weber, I. Zweyer, J. Wagner, C.E. Elger, E. U. Weber, E. J. Johnson

**Author notes:** Equally contributing first authors, listed in alphabetical order (A. Z. Enkavi, B. Weber).

## Abstract

Consistent decisions are intuitively desirable and theoretically important for utility maximization. Neuroeconomics has established the neurobiological substrate of value representation, but brain regions that provide input to the value-processing network is less explored. The constructed-preference tradition within behavioral decision research gives a critical role to cognitive processes that rely on associations, suggesting a role for the hippocampus in making decisions and to do so consistently. We compared the performance of 31 patients with mediotemporal lobe (MTL) epilepsy and hippocampal lesions, 30 patients with extratemporal lobe epilepsy, and 30 healthy controls on two tasks: binary choices between candy bars based on their preferences and a number-comparison control task where the larger number is chosen. MTL patients make more inconsistent choices than the other two groups for the value-based choice but not the number-comparison task. These inconsistencies increase with the volume of compromised hippocampal tissue. These results suggest a critical involvement of the MTL in preference construction and value-based choices.

**Significance:** Our days are full of choices that reflect our preferences. Economics lays out models of how to optimally make these decisions. Neuroeconomics has identified a cortical value-processing network whose activity correlates with constructs related to valuation and choice in economic models. However open questions remain: How are these value signals formed, and what regions might be necessary for retrieving and computing these value signals? Inspired by cognitive models calling on associative processes in value-based decisions, this paper uses unique neuropsychological data to establish the critical role of the medial temporal lobe in making consistent choices and further informs our understanding of the value-processing network.

## Introduction

Decision neuroscience made significant progress in identifying neurobiological correlates of value representations using paradigms involving simple choices between two stimuli based on underlying preferences (Hare, Camerer, & Rangel, 2009; Plassmann, O’Doherty, & Rangel, 2007). A value network involving a fronto-striatal circuit including the ventral striatum (VS) and the ventromedial prefrontal cortex (vmPFC), and posterior cingulate cortex (PCC) has been proposed (Bartra, McGuire, & Kable, 2013; Haber & Knutson, 2010). An unsolved question is where the value signals processed by this network come from, particularly for complex stimuli.

One influential conceptualization of preference construction proposes multiple steps including retrieval of relevant experiences with stimuli in the choice set, comparison of relevant attributes to reach a decision value, imagining future consequences of potential choices, that can be categorized as memory-related processes (retrospective or prospective) (Rangel, Camerer, & Montague, 2008; Weber and Johnson, 2009)

A long line of work in cognitive neuroscience shows the importance of the medial temporal lobe (MTL) in these processes (Squire, Stark, & Clark, 2004). The involvement and interaction of the MTL with the value network only recently attracted attention (Shadlen and Shohamy, 2016). Wimmer and Shohamy (2012) show MTL involvement in the value transfer of rewarded stimuli by associative learning that biases later decisions on non-rewarded stimuli. Barron, Dolan, and Behrens (2013) show activity in the hippocampus, in addition to medial prefrontal cortex, when subjects were asked to indicate preferences for novel food items based on familiar, but previously uncombined tastes. Gluth et al. (2015) show that choices are limited by memory constraints, which is associated with functional connectivity between the hippocampus and vmPFC (Gluth, Sommer, Rieskamp, & Büchel, 2015). Work motivated by the hippocampus’ involvement in imagining future experiences (Hassabis, Kumaran, Vann, & Maguire, 2007; Schacter, Addis, & Buckner, 2007) find that participants asked to imagine future events make more patient value-related decisions across time, which correlates with stronger activity in a set of brain regions including the hippocampus (Peters & Büchel, 2010). Impairment of these structures relates to more impatient choices, as shown in patients with subjective cognitive impairments regarded as a pre-stage of neurodegenerative disorders (Hu et al., 2017).

These studies suggest the involvement of the hippocampus and memory processes in value-related decision-making, but do not provide conclusive evidence that these processes are needed for such decisions. Such evidence requires comparing value-related decision-making abilities in the absence or impairment of these brain regions. Finding such differences would substantiate psychological models of decision-making involving memory processes and extend our understanding of the neural value network and the origins of value signals for complex options. Work that established the role of the ventromedial frontal region as crucial in the value network used this method: Patients with damage in these areas performed poorly in value-related decisions compared both to healthy controls, as well as patients with lesions elsewhere in the frontal cortex (Camille, Griffiths, Vo, Fellows, & Kable, 2011; Fellows & Farah, 2007).

Given these findings, we ask whether patients with hippocampal sclerosis are impaired in making consistent value-based decisions. Hippocampal sclerosis is a key neuropathological feature in patients with mesial temporal lobe epilepsy (Berkovic et al. 1991), with neurosurgical removal of the medial temporal lobe showing a high seizure-free rate. These patients show neuropsychological deficits mainly in the memory domain (Lin, Mula & Hermann, 2012; Hoppe, Elger, & Helmstaedter, 2007). To control for other epilepsy-related factors, like anticonvulsive medication or social effects of having seizure, we included in addition to healthy controls, a control group of patients with lesions outside of the temporal lobe. We test the affection of value-based decisions with binary choices among familiar food products. Our measure of choice quality is transitivity, the degree to which preferences are internally consistent. If a person chooses (Fig. 1.) Rolo over Bounty, and Bounty over Mars, choice transitivity requires they pick Rolo over Mars (Samuelson, 1938). Decision neuroscience uses this metric to quantify choice quality (Camille et al., 2011; Fellows & Farah, 2007; Fellows, 2006a; Kalenscher, Tobler, Huijbers, Daselaar, & Pennartz, 2010). As in this previous research, we included a pairwise judgment (rather than preference) task as a control, presenting respondents with pairs of numbers and asking them to judge which of the two is larger. This protocol is similar to that used to establish the necessary role of the vmPFC in value-related decisions (Fellows & Farah, 2007). Thus, selective differences in patients with MTL damage in value-based choices compared to numerical decisions should provide strong evidence for the involvement of the hippocampus, and thereby mnemonic processes, in value-based decision-making.

**Fig. 1.**
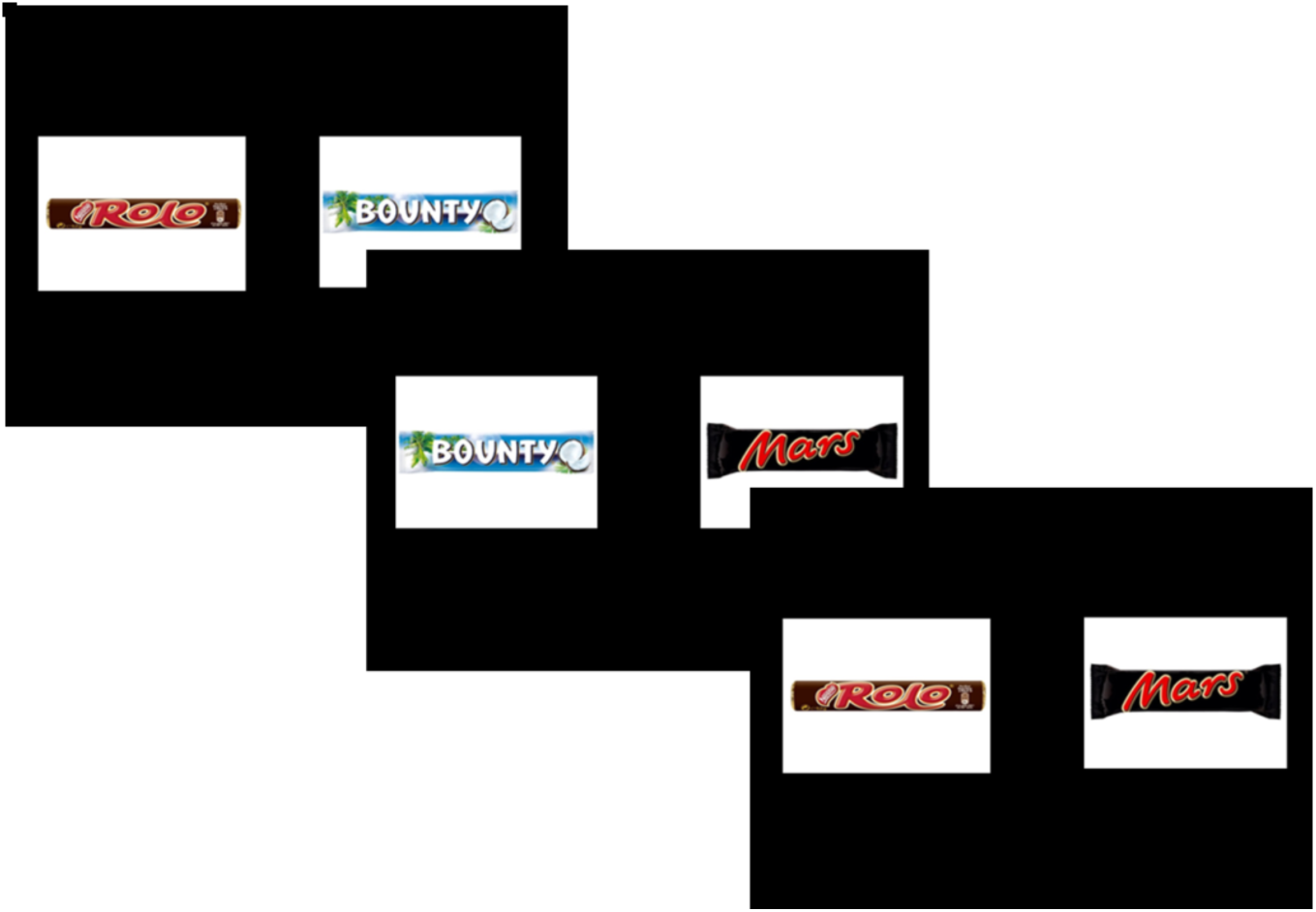
Three trials of the binary choice experiment. Subject indicated their preferred candy bar on each trial. Stimulus presentation and choice was self-paced, with a maximum length of 5 seconds.

## Methods

The study was approved by the local ethics committee of the University of Bonn and the Institutional Review Board at Columbia University (IRB-AAAB1301) and all subjects gave their written informed consent.

A total of 91 respondents participated. Thirty-one patients (15 female; mean age 47.74 with SD 2.56) suffering from mesial temporal lobe epilepsy with clinically diagnosed unilateral (left:n=14;right:n=8) or bilateral (n=9) hippocampal sclerosis from the presurgical program at the Department of Epileptology in Bonn were included in the study (here on referred to as MTL group). Different from patients with lesions in the vmPFC (Fellows & Farrah, 2007), the lesion locations in MTL patients are very similar. This makes lesion volume a better individual difference marker, as described below. Two control groups consisted of thirty patients with extratemporal lobe epilepsy (14 female; mean age 43.10 with SD 2.60; ETL group) and thirty healthy control subjects (15 female; mean age 51.40 with SD 2.60; CON group), respectively.

Each respondent made a series of choices between pairs of 20 candy bars, presented pictorially on a computer as in Figure 1. Each pairwise combination was presented once, resulting in (20x19)/2 = 190 choices for each participant, with a different random order. In a control task, subjects were presented with pairs of numbers, drawn from the range of one to twenty, and had to judge which number was larger. We computed judgment inconsistency across triplets of comparison identically for the two tasks. Subjects knew that they would receive their candy bar of choice from one randomly selected choice trial, in addition to a participation fee of 10 €.

Our focal dependent measure was the proportion of intransitive choices. A triplet is intransitive if (i) A was chosen over B and B was chosen over C, yet C was chosen over A or (ii) if B was chosen over A and C was chosen over B, yet A was chosen over C. (Fig 1. E.g. A can be Rolo, B, Bounty and C, Mars as described in the Introduction).

The proportion of intransitive choices was obtained by dividing the number of intransitive triplets by the total number of triplets. Analytically, it can be shown that the maximum level of intransitivities (those produced by a random responder) is 25% of all triplets. Below we report the results of simulations that demonstrate the non-linear relationship between number of intransitive choices and response error.

We also obtained, for a random subgroup of the patients with unilateral hippocampal sclerosis (n=16), a 3D-T1 weighted high-resolution data set (MP-RAGE, voxel size 1x1x1mm, repetition time 1570ms, echo time 3.42ms, flip angle 15°, field of view 256mm × 256mm) for volumetric measurement of the hippocampus. This was done in a fully automated manner by means of the FreeSurfer image analysis suite (Version 5.1.0, Martinos Center, Harvard University, Boston, MA, USA.; FreeSurfer, RRID:SCR_001847) (Fischl et al., 2002, 2004). Because of the high variance in total hippocampal volume between individuals, we used a lateral damage index of hippocampal volume to express the extent of unilateral hippocampal damage in our MTL group:

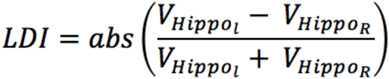

This lateral damage index can obviously be only assessed for subjects with unilateral hippocampal sclerosis.

### Experimental Design and Statistical Analysis

Our sample size was constrained by the availability of MTL patients with the appropriate lesion. Still, a power analysis based on effect sizes for healthy participants in the literature suggested we were well powered. Assuming a base proportion of 3% of intransitivities for healthy controls (Lee, Amir, & Ariely, 2009) and the same for ETL patients in contrast to twice this amount for the MTL patients, a large (and therefore conservative) estimate (i.e. an effect size of f = 0.4), we would need a total of 60 participants for a power level of 0.95. Our sample with at least 30 subjects per group was well above this.

To perform statistical analysis on our focal behavioral dependent measure, the intransitivity proportions were log transformed to avoid non-normal distributions and unequal variances between the tasks. Based on model comparisons, a linear mixed model was deemed the appropriate analysis having compared it to simpler models with no random effects. The contrasts of this model were orthogonalized to allow a direct comparison of the ETL group to the healthy controls and of the MTL group to both control groups together.

Statistical analyses were performed using R (Version 3.3.2; R Project for Statistical Computing, RRID:SCR_001905) for Mac. We use a two-tailed p-value of 0.05 as our criterion for statistical significance. The details of multilevel models are reported in the Results section.

## Results

As shown in Figure 2, MTL patients showed a greater percentage of intransitive choices compared to the two control groups in the preference task, but not in the control task (mean percentages for the preference task: MTL: 6.07%; ETL: 3.37%; CON: 2.75%; median percentages: MTL: 4.56%; ETL 2.72%; CON: 2.94%; mean percentages for the control task: MTL: 0.50 %; ETL: 1.00%; CON: 0.14%, median percentages: MTL: 0.36%; ETL: 0.00%; CON: 0.04%. This analysis used a linear mixed model regressing log transformed intransitivity percentages on an interactive model of group and task factors with orthogonal contrasts. The MTL-group task interaction was b = – 0.06, t(91) = –2.98, p = 0.004). The difference between degree of intransitivity between the preference and control task did not differ significantly between the two control groups (linear mixed model with orthogonal contrasts ETL-group task interaction b = – 0.04, t(91) = 0.97, p = 0.333).

**Fig. 2.**
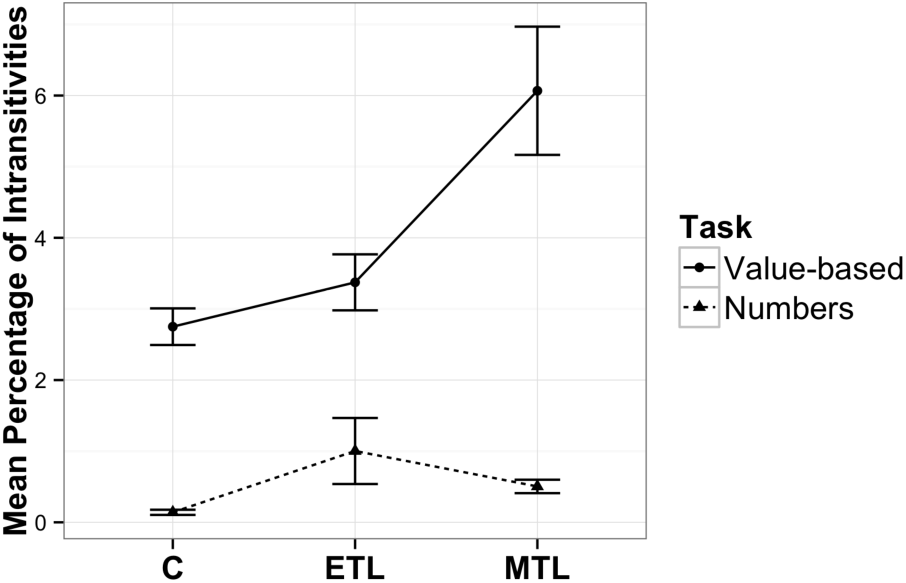
Mean percentage of intransitive choices per group in each task (n_MTL_ = 31, n_C_ = 30, n_ETL_ = 30). Error bars represent SEM.

For a subset of patients with available MRIs we determined the ratio of compromised hippocampal volume to total volume and correlated this individual difference variable with the percentage of intransitive choices observed for these participants. We used a non-parametric correlation coefficient that is insensitive to outliers because it is calculated using rank order. We found a strong and significant relationship between these two variables, as shown in Figure 3 (Spearman-rho = 0.676; F(1, 14) = 11.78, p=0.004; n=16), such that the larger the lesion volume, the less consistent were the value-based choices.

**Fig. 3.**
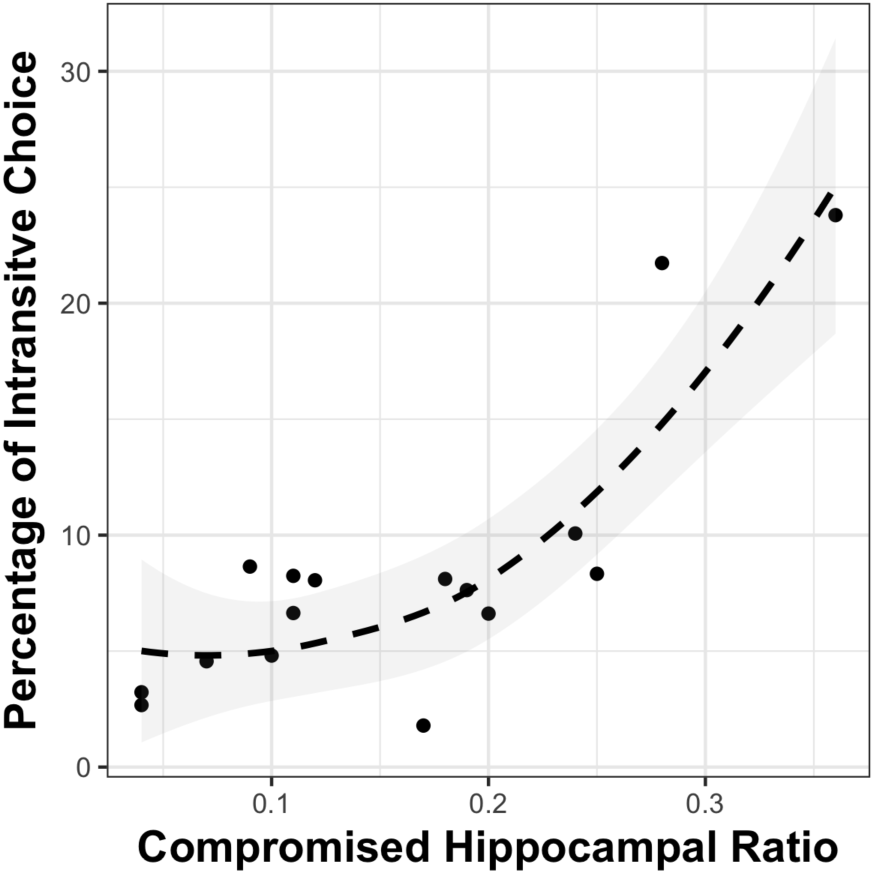
Relationship between hippocampal lesion volume and intransitive choices. Scatterplot of compromised hippocampal volume (as a ratio of total volume) against percentage of intransitive choices. Smoothing is done with locally with α = 2. The observed robust nonparametric rank order correlation rho=0.676, p=0.004.

To provide context for interpreting the observed frequencies of intransitivity, we conducted a series of simulations that use a random utility model with a stochastic term added to the utility of the options, such that the probability of choosing option A(*p*(*A*)) in a decision between A and B is:

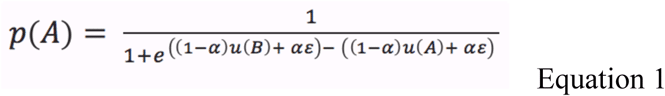

where *u(A)* and *u(B)* represent the utilities of options *A* and *B*, α represents the proportion (between 0 and 1) of the observed utility due to random error, and ε is the random error. It can be shown analytically that the maximum proportion of intransitive triples is .25 (Figure 4, also see the discussion section of Tversky, 1969). Our question of interest is the effect of α, the proportion of random error upon intransitivity. Our hypothesis is that the degree of MTL patients’ hippocampal sclerosis increases α, since access to past experiences that would normally be called on to make a choice is impaired. We simulated how the proportion of intransitive triples increases as noise in utilities increases. The effect is non-linear (Figure 5), and the observed intransitivities in the MTL group correspond to an α of .3, i.e., the level expected if random error represented approximately 30 percent of the utility values in Equation 1.

**Figure 4:**
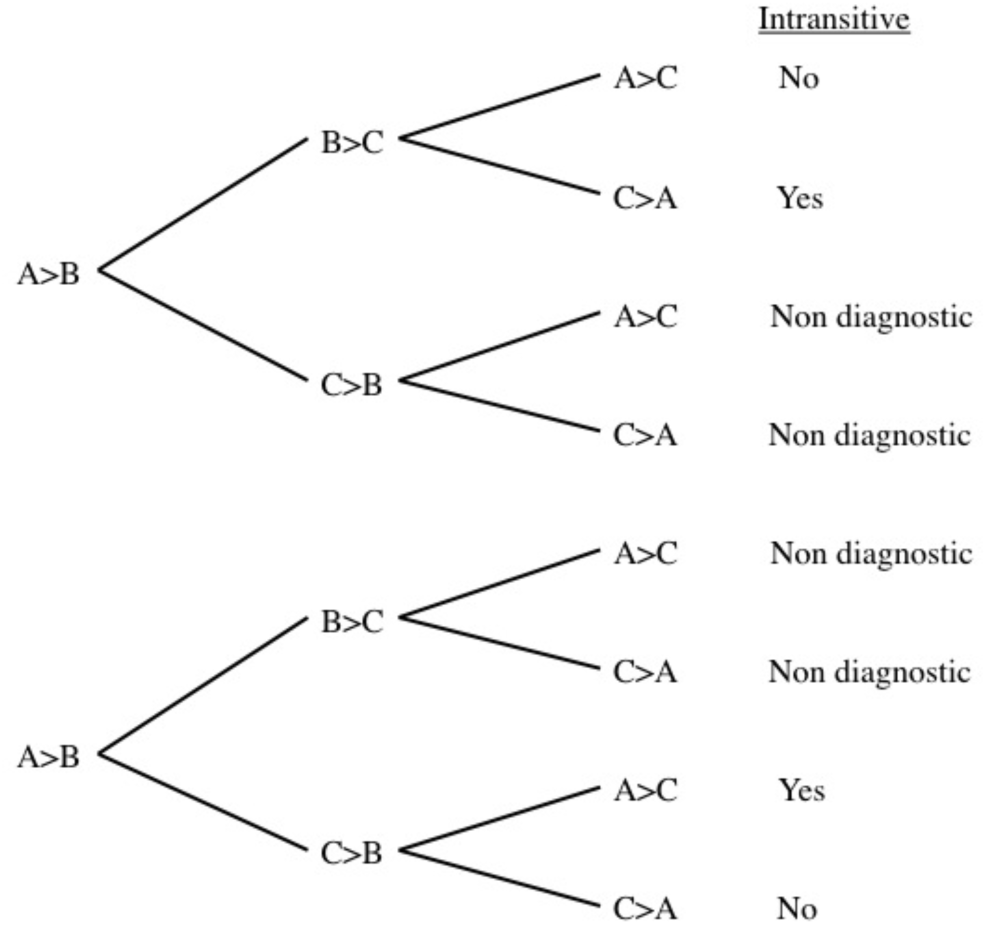
Tree diagram indicating possible intransitive paths from three binary choices.

**Figure 5:**
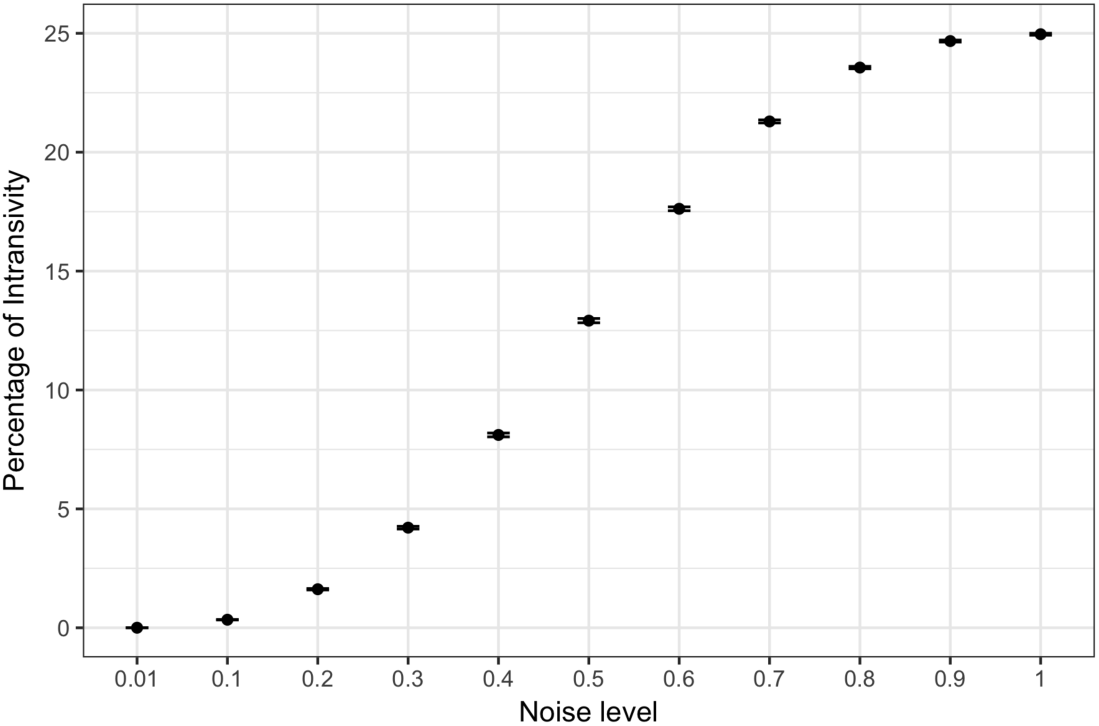
Mean percentage of intransitivities at different noise levels (α), based on 1000 simulations at each noise level. Error bars indicate standard errors.

We can test several alternative explanations to our account of random error in value construction for our data. One alternative explanation is that respondents retain explicit episodic memory of previous value comparisons during the task, and do not perform value construction for the two options of each pairwise choice. Under this account, non-MTL respondents may have better memory for their choices made earlier in the task, and this better episodic memory prevents intransitive choices. This account would suggest that the rate of intransitivities declines over time, as previous choices are remembered and used to avoid intransitive later choices. We might expect this decline in intransitivities over choice trials would differ for the MTL and non-MTL groups. We tested this hypothesis by regressing whether or not a triplet was intransitive on the trial number of the last seen trial in that triplet. We found no increase in the probability of a triplet being intransitive depending on when the subjects saw the last trial in that triplet (b = 0.027, z = 0.79, p = 0.427) nor was this trend different for the MTL group (b = 0.032, z = −0.78, p = 0.434).

An account emphasizing episodic memories of previous choices during the task makes a more specific hypothesis: It would predict that the probability of instransitivity depends on the delay (number of trials) between the choices involving the items that define an intransitive triplet. To test this we checked whether a triplet was more likely to be intransitive depending on the variance in the trial numbers involved in that triplet. We found that the further apart from each other the three choices in a triplet were made the more likely they were to be intransitive (b = 0.109, z = 3.40, p = 0.007). Crucially, however, this pattern was not different for the MTL group (b = −0.049, z = 1.25, p = 0.213). That is, the group differences in intransivity cannot be explained by impairments of episodic memories *during* the task.

Another alternative explanation involves group differences in speed-accuracy tradeoff. To test this, we examined response latencies of the choices, and the relationship between responses latencies and intransitivities for MTL and non-MTL groups. Contrary to a speed-accuracy tradeoff, we found that slower (rather than faster) trials were more likely to be involved in intransitive triplets (b = 0.441, t(16985) = 4.40, p = 0.00001) for all groups, and that this did not differ for the MTL group (i.e., no interaction with this group: b = −0.0846, t(16985) = −0.62, p = 0.535, though there was a quadratic effect of time for the ETL group b = −0.382, t(16985) = −2.69, p = 0.007). Moreover, the MTL group actually had a significantly slower average response time per trial (b = 0.301, t(88) = 2.11, p = 0.038). Together, these results suggest that intransitive triplets accompany more effortful and longer responding, eliminating the possibility of a speed-accuracy tradeoff.

Notably both the speed accuracy tradeoff and the effect of the trial number of the last trial in a triplet on intransitivity is the same for the numbers task as it is for the choice task, suggesting that the two tasks share some similarities. Finally, we examined whether there were any idiosyncratic effects on preference intransitivity associated with specific stimuli (candy bars). We found no significant differences in the average number of intransitive triplets each candy bar was involved in (F(1, 90) = 0.003, p = 0.955).

In combination, these analyses suggest that the observed increase in transitivity violations for respondents with MTL lesions in the preference task but not number-comparison task, in a way that is related to the volume of hippocampal lesions, suggests a failure in value-related associations in this group.

## Discussion

We provide support that brain regions associated with memory-related associative processes play a critical role in value-based decision-making. Hippocampal lesions are associated with an increase in intransitive value-based choices, and the degree of intransitivity is related to magnitude of the damage to the hippocampus. A control task not involving value-based processes does not show these effects, nor do respondents who have lesions outside of the medial temporal lobe. These dissociation results implicate a crucial role for the hippocampal areas in preference construction (Lichtenstein & Slovic, 2006), a conceptualization in behavioral decision research that contrasts with standard theories of rational choice that implicitly assume stable utility functions and choice options with preexisting values.

Two conceptual clarifications are in order. Our central dependent measure, the frequency of intransitive preferences has been used before to examine the inability of decision makers to produce a stable representation of the value of choice options, with other patient groups (Camille et al., 2011; Fellows & Farah, 2007). Earlier work, however, using choice intransitivity as a dependent measure did so to identify choice heuristics incompatible with utility maximization (Tversky, 1969). This resulted in a debate on the correct probabilistic model of transitivity that would account for errors in experimental data and whether that was evidence for a particular mechanism (Birnbaum & Gutierrez, 2007; Regenwetter, Dana, Davis-Stober, & Guo, 2011; Regenwetter & Davis-Stober, 2008). Our use of the term pairwise “transitivity” is not based on these frameworks and our design with two alternatives per choice does not employ such model comparison. We use intransitivity counts, as in other decision neuroscience research, instead, to examine error associated with the construction of value representations.

Second, our use of the term “transitivity” is only marginally related to the extensive literature measuring transitive inference, where a set of premises are learned in the experiment and participants are asked to generalize these learned rules to novel contexts and combinations of stimuli. Transitive inference tasks have been instrumental in establishing the role of the hippocampus in representing organizations of stimulus relations (Eichenbaum & Cohen, 2001). Animal lesion studies established the necessity of the hippocampus for transitive inference (Bunsey & Eichenbaum, 1996; Dusek & Eichenbaum, 1997), and data from humans has confirmed the involvement of this region (Heckers, Zalesak, Weiss, Ditman, & Titone, 2004; Nagode & Pardo, 2002). However transitive inference paradigms differ from ours, critically, because our respondents are stating their preferences, not learned premises. We do not present participants with transitive relations and ask them to reason following this rule. We ask for their preference between two candy bars. We do *not* hypothesize that if a participant chooses Snickers over Mars and Mars over Bounty they would also choose Snickers over Bounty because they are instructed that these choices must follow a given transitive relationship. Instead, their transitive choice reflects an anticipation that they will enjoy Snickers more. That is, while a transitive inference task implies a strict ordinal relationship between stimuli thereby recruiting working memory, transitivity of choice as measured by our design relies on values learned over time and presumably relies on the recruitment of associative facilities (Halford, 2005).

Despite the evidence for the involvement of the hippocampus in consistent value-based decisions, the delineation of specific cognitive and neural mechanisms provide multiple avenues for future research.

First, the hippocampus is just one part in a larger network of relevant brain areas involved in the retrieval and processing of choice values. A recent review (Shohamy & Turk-Browne, 2013) suggests hippocampal involvement in a variety of cognitive functions outside of the domain of declarative memory providing two different hypotheses of hippocampal function: The memory modulation hypothesis proposes that representations within the hippocampus may transiently bias other cognitive functions e.g. value computations in our task. The adaptive function hypothesis, in contrast, highlights the hippocampus as a central processing unit with specific computations carried out in the hippocampal networks, depending on the task at hand.

Our hippocampal patients produce patterns of intransitivity of value-based choice that are similar to those observed in ventromedial prefrontal cortex (vmPFC) patients, suggesting that the associations and memories stored in the hippocampus may serve as inputs to value calculation occurring elsewhere (Barron et al., 2013), potentially in line with the memory modulation hypothesis. The hippocampus is one of the most highly interconnected brain areas (Cole, Pathak, & Schneider, 2010; Godsil, Kiss, Spedding, & Jay, 2013). In addition to being directly and monosynaptically connected to the prefrontal cortex, animal work suggests a topographically specific hippocampal projections map on functionally distinct prefrontal regions (Cole et al., 2010; Godsil et al., 2013).

This possibility calls for a nuanced investigation of the interactions between hippocampal and prefrontal regions in value-based decision-making. For example, Ranganath and Ritchey (2012) propose a division of the MTL into two systems for memory-guided behavior: the anterior (AT) and posterior-medial (PM) system. The AT, which is comprised of the perirhinal cortex and anterior parts of the hippocampus and amygdala has strong interconnections with the frontal cortex, has been argued to be involved in familiarity-based cognition, social behavior and saliency. This is also the part of the hippocampus which is most affected in patients with hippocampal sclerosis (Woermann, Barker, Birnie, Meencke, & Duncan, 1998). Ranganath & Ritchey (2012) suggest that the AT system could facilitate the use of past experiences to inform inferences about the personality and intentions of others. Our results suggest such inferential abilities specific to distinct regions in the MTL along with the connection to the ventromedial prefrontal cortex may play a role in value-based decisions.

On the other hand, in line with an adaptive function hypothesis, deficits in consistent choices might be due to hippocampus-specific computations. For example, Fellows, (2006b) showed that vmPFC lesioned patients differ from normal controls in their external information search, in ways that could be attributed to diminished planning capacity. Perhaps this planning capacity relies on hippocampus-specific computations. An interesting topic of research would be whether vmPFC patients exhibit deficits in different mnemonic processes.

A second future research topic are potential compensation mechanisms in patients with chronic hippocampal lesions. It is well-known that chronic brain lesions may lead to compensatory shifts in neural processes, e.g. in the domain of language processing (Weber et al., 2006). The application of neuroimaging methods during a value-based decision task in these patients could provide answers to this question.

Third, although patients with temporal lobe epilepsy and hippocampal sclerosis do show neuropsychological deficits especially in the domain of declarative memory, the amount to which these deficits occur varies strongly between patients (Hoppe, Elger, & Helmstaedter, 2007). Future research combining in-depth neuropsychological testing together with value-based choice tasks may shed light on the specific cognitive components underlying the observed range of decision deficits.

Our results suggest a critical role for the hippocampus in the construction of the value of choice options. Most decisions require the construction of value based on past experience. Even a previously experienced option, like a favorite dish in a familiar restaurant, requires us to compare recollections of the value of that option to newly available options such as tonight’s specials. A better understanding of both internal and external inputs to preference construction processes and their aggregation and comparison will allow us to comprehend and model how the brain calculates value and makes consistent choices.

## Acknowledgements

BW was funded by a Heisenberg-Grant of the German Research Council (WE 4427/3-2) and EUW and EJJ by NIA Grant 5R01AG027934. AZE is currently at the Department of Psychology of Stanford University and EUW is at Woodrow Wilson School, Andlinger Center for Energy & Environment, and Department of Psychology, Princeton University. The authors declare no competing financial interests.

## Author contribution statements

BW, EJJ and EUW designed the experiment and wrote the manuscript, EJJ and AZE analyzed the behavioral data and wrote the manuscript, IZ performed experiments, JW analyzed the MRI data. CEE provided clinical data of the patients. All authors approved the final version of the manuscript for submission.

